# Trustworthy detection of exencephaly in high-throughput micro-CT embryo screens with focal-loss transformers

**DOI:** 10.1101/2025.08.12.669840

**Authors:** Oshane O. Thomas, Rachel Roston, Hongyu Shen, A. Murat Maga

**Affiliations:** Center for Development Biology and Regenerative Medicine, Seattle Children’s Research Institute, Seattle, Washington, United States of America; Amazon.com, Seattle, WA, USA; Division of Craniofacial Medicine, Department of Pediatrics, University of Washington, Seattle, Washington, United States of America

## Abstract

Lethal and sub-viable knockout mouse lines require whole-embryo 3D imaging to connect genotype to phenotype (Dickinson et al., 2016; Cacheiro et al., 2022). There are often far fewer samples of in-class (e.g., homozygous knockouts) than wildtype or normative samples. Such extreme subject-level imbalance degrades both statistical anatomy and deep learning, often yielding saliency maps that highlight noise rather than lesion-specific signal (Adebayo et al., 2018; Buda et al., 2018; Johnson & Khoshgoftaar, 2019). We therefore asked whether focal loss (Lin et al., 2017) in combination with model-capacity control and seed ensembling can stabilize explanations without compromising classification accuracy.

Exencephaly is an neural tube defect characterized by incomplete closure of the cephalic neural tube and dorsally exposed,disorganized neural tissue (Greene & Copp, 2014; Noden & de Lahunta, 1985). In our late-gestation screen, approximately 10% of embryos had exencephaly, creating severe class imbalance and underscoring the need for interpretable automation.

As a test case for imbalance-aware, interpretable phenotyping, we analyzed 253 diceCT scans of E15.5 embryos (24 with exencephaly). A self-supervised transformer was fine-tuned with three regimes: cross-entropy (CE-Large), focal-loss equal-capacity (FL-Large) and focal-loss reduced-capacity (FL-Small). Five random seeds per regime yielded 15 models. Integrated Gradients saliency was quantified, and explanation quality was measured by saliency entropy (sparsity), cross-seed Dice/Jaccard similarity (reproducibility), and expert visual inspection.

All 15 models achieved near perfect phenotype recognition on held-out data with 0.996 ± 0.002 mean accuracy with some seeds/regimens reached 1.000. Focal loss reduced saliency entropy by up to 1.5 bits and doubled cross-seed Dice overlap, concentrating attribution on the malformed cranial vault. Ensemble-averaged heat-maps show that both focal-loss regimes concentrate attribution on the malformed cranial vault while suppressing spurious body-wide signals.

Focal loss, modest capacity, and seed ensembling within a modified M3T transformer yielded sparse, reproducible, anatomically focused attribution while preserving perfect sensitivity. This supports trustworthy high-throughput phenotyping in severely imbalanced embryo screens. The workflow relies only on standard atlas registration and image pre-processing, requires no voxel-level annotations, and is readily adaptable to other structural malformations and developmental stages.

## 3. Introduction

Thousands of genes are essential for mammalian development; when disrupted, embryos often fail to survive, leaving prenatal imaging as the primary window into gene function for lethal and sub-viable lines (Dickinson et al., 2016; Cacheiro et al., 2020). In large-scale knockout programmes such as the International Mouse Phenotyping Consortium (IMPC) and the NIH Knockout Mouse Phenotyping project (KOMP), late-gestation embryos are routinely imaged to capture whole-embryo morphology and viability because many lethal strains cannot be evaluated postnatally (Dickinson et al., 2016). These pipelines commonly acquire iodine-contrast micro-CT (diceCT) volumes in late gestation (e.g., E15.5) that resolve both soft tissue and mineralising bone in a single scan and provide standardized 3D readouts suitable for cross-line comparison (Wong et al., 2013; Dickinson et al., 2016). The growing abundance of these high-resolution 3D image datasets require high-throughput computational methods to rapidly detect and characterize morphological phenotypes associated with genetic mutations. Multiple proposed pipelines use image registration-based approaches to detect subtle, quantitative phenotypic differences (i.e., subtle differences in size and shape of anatomical structures) (e.g., Dickinson et al., 2016; Horner et al., 2021; Wong et al., 2014). Often, these approaches are unsuitable for the detection of qualitative or overt phenotypes, such as large-magnitude quantitative differences or missing structures, that are common in mutant screens (Dickinson et al., 2016; Groza et al., 2023; Horner et al., 2021).

Deep learning-based classification offers a potential approach to detect and characterize qualitative phenotypes in large, high-throughput genetic screens. However, class imbalances of subjects and small subject sample sizes with a relatively large number of voxels per image result in a “double disparity”: abundant, highly correlated spatial features but few independent observations.The number of subjects with the phenotype of interest is often relatively small compared to the number of unaffected subjects, particularly when mutations are subviable or lethal, and the morphology of the structural defect is also often highly variable because of incomplete penetrance and variable expressivity (Horner et al. 2021). Furthermore, the phenotype of interest often comprises a relatively small percentage of the total number of voxels in an image. High-resolution micro-CT and micro-MR images of late-gestation embryos typically have isotropic spacing of 18–50 µm to capture small defects in organogenesis (Wong et al., 2013). At these resolutions a single whole-body image yields on the order of 10⁷– 10⁸ voxels, while structural defects within an organ comprise a much smaller number of voxels. If each voxel is considered a single measurement, a research study would contain orders of magnitude more measurements than subjects. This combination undermines assumptions of independence that underlie classical voxel-wise inference and mini-batch optimisation in deep learning (Ioannidis, 2005; Litjens et al., 2017). Practically, millions of voxel-wise comparisons across a few dozen embryos inflate false-discovery rates and erode statistical power, while class imbalance drives naïve learners toward the majority class and yields deceptively high accuracy with poor sensitivity to the minority phenotype (Ioannidis, 2005; Johnson & Khoshgoftaar, 2019). Deep networks add a second fragility: gradient-based attribution methods can appear plausible yet remain insensitive to trained parameters, reflecting input edges rather than learned pathology, especially under overfitting (Adebayo et al., 2018). For high-throughput embryo phenotyping, where explanations guide downstream histology and hypothesis generation, attribution must be anatomically specific and reproducible.

Here, we aim to address two linked requirements of using a deep learning-based approach for phenotype classification and characterization : reliable detection of relatively rare malformations and the generation of anatomically meaningful explanations. Our approach embeds three safeguards in a self-supervised transformer framework and defers full details to Materials and Methods. First, class-balanced mini-batch sampling equalises class exposure during optimisation so that gradients are not dominated by normal embryos (Buda et al., 2018). Second, focal loss reshapes the objective to down-weight easy negatives and preserve gradient signal on rare positives; in our setting, voxels corresponding to the malformed cranial vault (Lin et al., 2020). Third, we limit model variance by reducing transformer depth and width, consistent with empirical scaling showing that excess capacity increases overfitting in small medical-image cohorts (Tan & Le, 2019). These measures act on different failure modes: data distribution, loss landscape, and hypothesis class. However, they share a common biological aim: to retain sensitivity to rare defects while suppressing spurious, body-wide explanations. To evaluate interpretability, we aggregate integrated gradient attribution across independently initialised models (seed ensembles) and register attribution to a common atlas, asking whether the resulting maps consistently highlight the cranial vault in affected embryos and remain neutral elsewhere (Lakshminarayanan et al., 2017).

In this study, we use exencephaly as a test case for our deep learning-based approach for phenotype classification. Exencephaly is a severe and lethal neural tube defect in which the cephalic neural tube fails to close completely, leading to disorganized neural tissue that remains externally exposed and not covered by the skin (Greene & Copp, 2014; Noden & LaHunta,1985) (Figure 1). In our E15.5 dataset, ∼10 % of embryos showed exencephaly, creating a subject-level imbalance that mirrors many biomedical detection problems in which the minority class is the phenotype of interest (Johnson & Khoshgoftaar, 2019). This relatively infrequent brain phenotype is also anatomically overt and spatially confined, enabling clear separation of signal from background and unambiguous expert verification.

**Figure 1.**
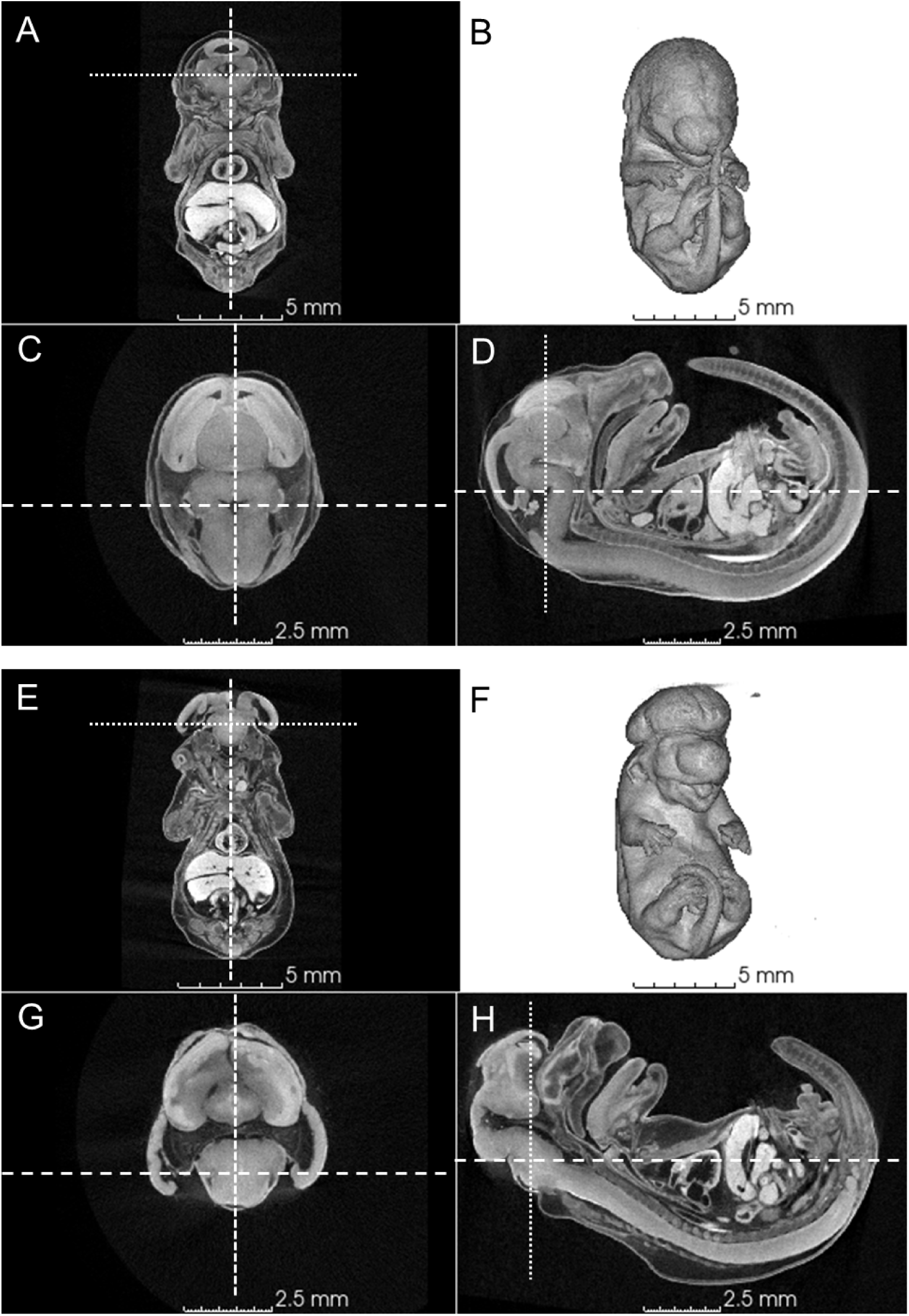
Representative diceCT volumes of a wild-type and an exencephalic E15.5 mouse embryo. Panels A–D depict a phenotypically normal (wild-type) embryo, and panels E–H depict an embryo with exencephaly. For each specimen, orthogonal cross-sections through the rigidly aligned volume are shown in coronal (A, E), axial (C, G), and sagittal (D, H) planes, with dashed lines indicating the locations of intersecting slice planes. Panels B and F show corresponding 3-D volume renderings. The wild-type embryo exhibits normal brain structures including the cerebral hemispheres, midbrain, brain stem, ventricles, and three flexures (A–D), whereas the exencephalic embryo shows three flexures but lacks normal anatomy of the dorsal structures due to failed closure of the neural tube (E–H). Voxel spacing = 18 µm³ for all images. Scale bars: 5 mm (B, F) and 2.5 mm (A, C, D, E, G, H).

Using a real E15.5 diceCT cohort with ∼10:1 normal-to-affected ratio, we preregistered three hypotheses: that replacing class-weighted cross-entropy with focal loss would reduce saliency dispersion (Hypothesis 1); that a capacity-controlled transformer would increase cross-seed agreement of saliency (Hypothesis 2); and that ensemble-averaged, atlas-registered Integrated Gradients would concentrate over the cranial vault in affected embryos and remain largely neutral in unaffected embryos (Hypothesis 3). The remainder of the paper reports tests of these hypotheses and examines how imbalance-aware training can support trustworthy, high-throughput embryo phenotyping.

## 4. Materials and Methods

Figure 2 provides an overview of the study pipeline from specimen harvest through using Integrated Gradients to obtain meaningful attribution maps.

**Figure 2.**
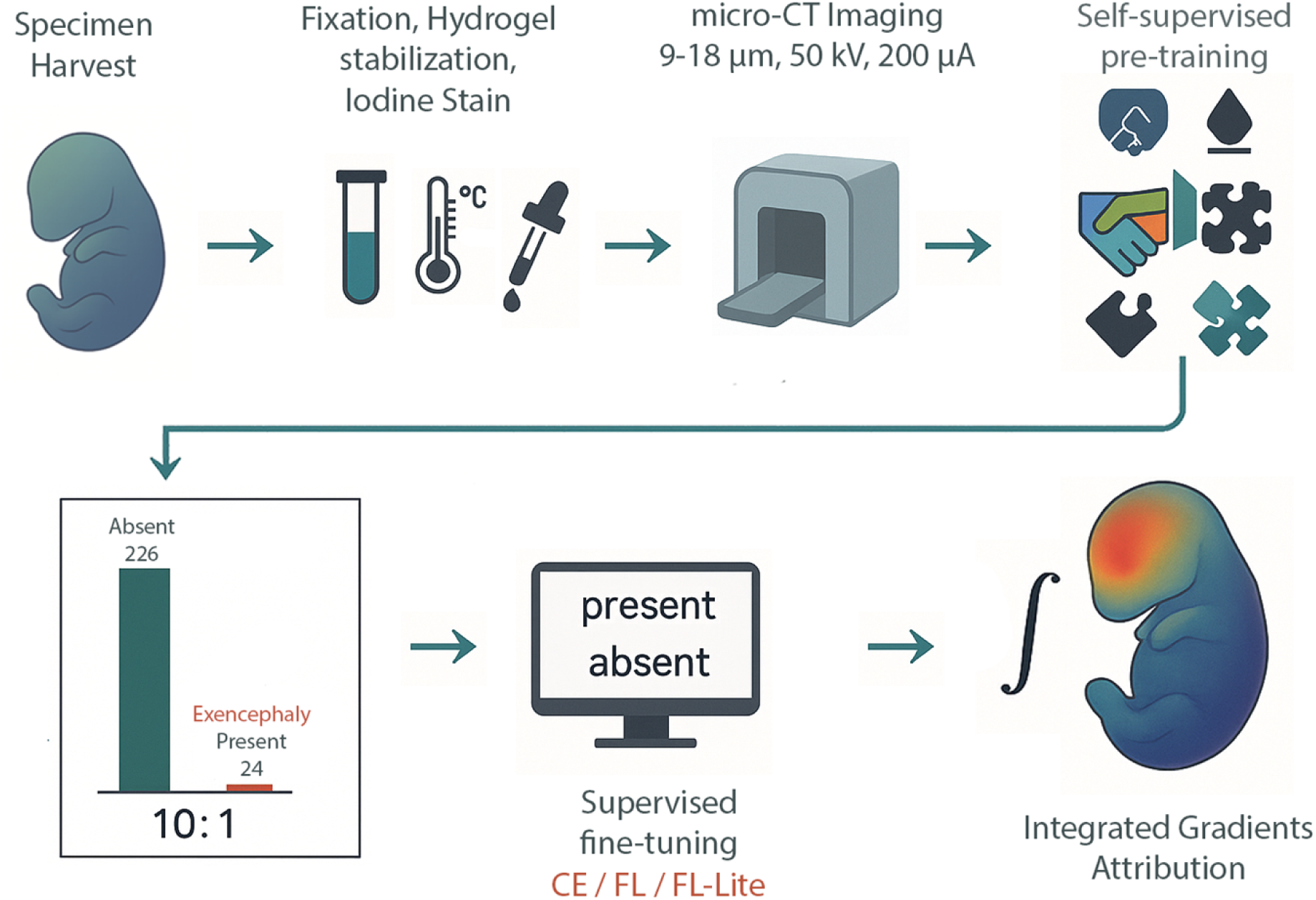
Study workflow and class imbalance in high-throughput embryo phenotyping. The pipeline begins with specimen harvest at embryonic day 15.5 (E15.5), followed by fixation in 4 % paraformaldehyde, hydrogel embedding, iodine contrast staining, and micro-CT imaging at 9–18 μm isotropic resolution (50 kV, 200 μA). The resulting dataset contains 250 total embryos, with a pronounced 10:1 class imbalance: 226 anatomically normal and 24 exhibiting exencephaly. These volumes undergo self-supervised pre-training using four proxy tasks, Bootstrap-Your-Own-Latent (BYOL), masked voxel reconstruction, 3D rotation prediction, and slice-order (jigsaw) classification, before supervised fine-tuning under one of three regimes: CE (cross-entropy baseline), FL (focal loss), or FL-Small (focal loss with reduced transformer capacity). Integrated Gradients (IG) attribution is then performed on correctly classified scans to identify voxels that most strongly influence predictions: seed ensembles and atlas-space registration yield stable, interpretable heat maps for downstream anatomical analysis.

### Dataset Description, Image Acquisition, and Preprocessing

All specimens were collected as part of an ongoing mouse reverse-genetic screen approved by the Seattle Children’s Research Institute Institutional Animal Care and Use Committee (IACUC 00030). Timed pregnancies from C57BL/6J matings were terminated at embryonic day 15.5 (E15.5), a stage widely used for lethality and morphology assessment in large-scale knockout phenotyping pipelines (Dickinson et al., 2016). The study cohort comprised 253 embryos: 24 with exencephaly and 229 without exencephaly. Embryos without exencephaly included phenotypically wildtype embryos as well as embryos with non-exencephaly structural defects (e.g., holoprosencephaly, craniofacial dysmorphologies, and overt differences in internal organs). Genotype information was not used at any stage because the objective was purely phenotypic classification (exencephaly present vs absent), consistent with image-based screening practice (Dickinson et al., 2016; Cacheiro et al., 2020).

Immediately after dissection, each embryo was immersion-fixed overnight in 4 % paraformaldehyde at 4 °C, washed in phosphate-buffered saline (PBS), and stabilised in a hydrogel matrix as described for embryo micro-CT phenotyping (Wong et al., 2013; Dickinson et al., 2016). Briefly, specimens were incubated at 4 °C for 72 h in hydrogel solution (4 % acrylamide, 0.25 % VA-044 thermal initiator, 0.05 % bis-acrylamide, PBS) and polymerised for 3 h at 37 °C (Wong et al., 2013). Soft-tissue contrast enhancement was achieved with Lugol’s iodine (0.1 N I2KI) for 24 h, followed by one rinse and two PBS washes, a standard diffusible iodine–based contrast-enhanced micro-CT (diceCT) approach for whole embryos (Metscher, 2009; Gignac et al., 2016). Finally, each embryo was embedded in 1 % agarose and scanned within 24 h on a Bruker SkyScan 1272 micro-CT system with a 1-mm aluminium filter. 3D scanning of embryos was conducted at the SCRI MicroCT Imaging Facility (RRID:SCR_024678), using Bruker Skyscan 1272 microCT that was funded by an NIH shared instrument grant (S10OD032302).

Where applicable, diceCT volumes acquired at 9 µm isotropic resolution were down-sampled to match the 18 µm native resolution of the remaining scans using the SkyscanReconImport batch workflow within SlicerMorph (Rolfe et al., 2021). Prior to analysis, each image was rigidly re-oriented to standard anatomical planes and down-sampled to 60 µm³ isotropic spacing to reduce memory requirements. To provide a fixed spatial canvas for learning, each volume was cropped with a 5-voxel margin around the embryo body and then resampled and padded to 144 × 144 × 144 voxels.

Intensities were converted to float and Z-score normalised using the global training-set mean (μ = 1,776.8) and standard deviation (σ = 5,603.5). The resulting dataset consists of intensity-normalised, orientation-standardised cubic volumes with uniform voxel spacing and dimensions, ready for the self-supervised pre-training and supervised fine-tuning described below. The case mix and overall class ratio approximate those encountered in late-gestation knockout screens, where embryo imaging is used to characterise lethal and subviable lines (Dickinson et al., 2016; Cacheiro et al., 2020; Gignac et al., 2016).

### Self-supervised pre-training to learn anatomy without phenotype labels

Because only 24 labelled positives are available, we first expose the network to ∼2 × 10⁷ unlabeled voxels per embryo so that it can internalise normal anatomical variation before seeing any phenotype labels. Goal: learn generic embryo anatomy from unlabelled data to reduce labelled-sample requirements. We used a single hybrid three-plane CNN–Transformer backbone for both self-supervision and downstream classification, ensuring that weights learned here are transferred intact to the fine-tuning stage. (Fig. 3) (Jang and Hwang 2022). A shallow 3-D stem (two 5 × 5 × 5 convolutions, 64 filters, dropout = 0.05) learns local context, after which the intermediate feature map is re-sliced into contiguous coronal, sagittal and axial stacks (3 × 144 = 432 images). Each slice is processed by a ResNet-50 whose fully-connected layer is replaced by a two-layer projector (either 128-D or 256-D, depending on the target embedding). Plane-specific and positional embeddings, together with a [CLS] token, form the input sequence for a pre-norm Transformer encoder (12 layers, 8 heads, 256-D; a reduced variant with 4 heads and 128-D is used later). DropPath is linearly increased from 0 to 0.05 within the self-supervision phase to regularise deep layers (Vaswani et al., 2017).

**Figure 3.**
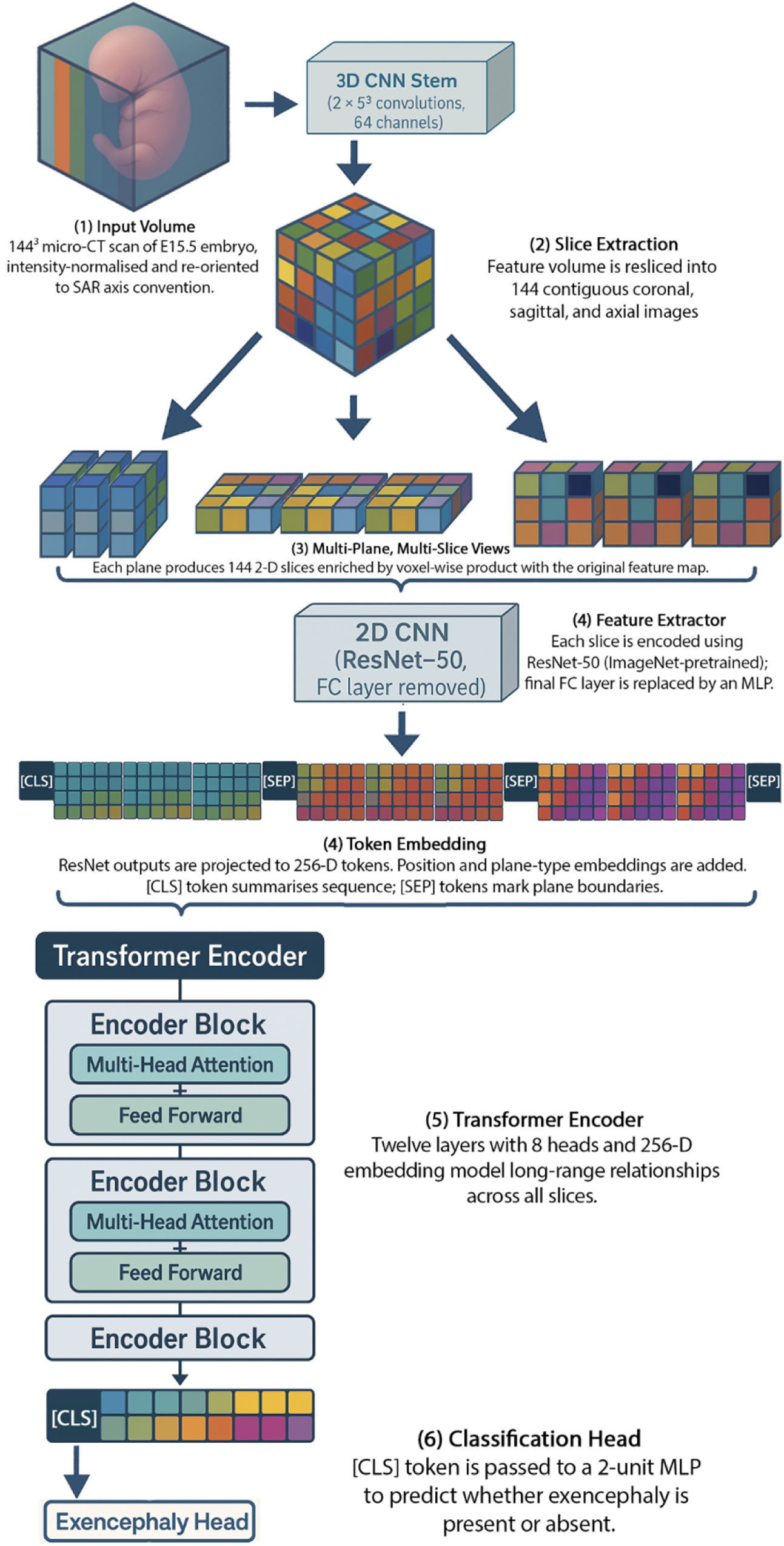
Hybrid three-plane transformer architecture for embryo classification. This model is adapted from the M3T architecture first introduced by Jang and Hwang (2022) for 3D medical image classification. A 144³ micro-CT volume is passed through a 3D CNN stem to capture local spatial features, then re-sliced into 144 contiguous images along each anatomical plane (coronal, sagittal, axial). A shared ResNet-50 processes each slice, and the resulting 2D features are projected to 256-dimensional tokens. These are interleaved with plane-specific, positional, and special tokens ([CLS], [SEP]) to form a 436-token input sequence. A 12-layer transformer encoder with 8 attention heads models long-range slice-to-slice relationships across the whole sequence (Vaswani et al., 2017). The final [CLS] token is decoded by a two-unit MLP to predict the presence or absence of exencephaly. This hybrid approach enables joint modelling of local 3D context and global planar dependencies within a unified transformer-based architecture.

#### Multi-objective self-supervision

To endow the encoder with general embryo anatomy while remaining agnostic to any downstream label, we combined four complementary self-supervised tasks into a single objective

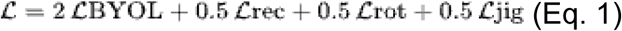

The weights (2:0.5:0.5:0.5) were selected by a small grid search on a linear-probe validation set and held fixed thereafter. Briefly, 𝓛_BYOL aligns latent representations of two independently augmented views to capture instance-level anatomy (Grill et al., 2020); 𝓛_rec reconstructs randomly masked voxels, forcing awareness of local tissue texture (He et al., 2022); 𝓛_rot predicts one of twelve right-angle 3-D rotations, promoting global spatial reasoning (Gidaris et al., 2018); and 𝓛_jig discriminates in-order from permuted slice stacks, sensitising the network to inter-slice continuity (Noroozi & Favaro, 2016). Full mathematical definitions are provided in Supplementary Methods, Eq. S1–S4. Together, the four losses expose the network to information ranging from single-voxel appearance to whole-brain orientation while requiring no phenotype labels.

#### Augmentation and schedule

For each volume two stochastic views are produced via MONAI 1.4 (Cardoso et al., 2022) transforms: random flips, Random Affine (±5°, ±5 %, ±4 voxels), contrast/scale/shift, Gaussian noise (σ ≤ 0.01), Gaussian smoothing (σ ≤ 1 voxel) and Random Coarse Dropout (four 16³ voids, p = 0.2) applied only to the second view. Volumes are resampled to 144³ voxels, Z-score normalised (μ = 1,776.8, σ = 5,603.5) and batched in twelves. Training proceeds for 800 epochs on a single NVIDIA A100-80 GB GPU (AdamW, lr = 6 × 10⁻⁴, weight decay = 3 × 10⁻⁶) with a 10-epoch linear warm-up followed by cosine decay (Zhuang et. al, 2022; ). Mixed-precision (FP16) with dynamic gradient scaling is used throughout. The run consumed ∼318 GPU-h; checkpoints were stored every ten epochs, and the final encoder weights are publicly released.

### Multi-plane multi-slice transformer classifier (M3T)

We modified and adapted the multi-plane-and-multi-slice transformer (M3T) initially developed for Alzheimer’s disease MRI classification and described in the previous self-supervised pre-training section to serve as a phenotype classifier for embryonic micro-CT (Jang and Hwang 2022) (Figure 3). All convolutional and transformer layers inherit the self-supervised weights described earlier. In our pipeline, the self-supervised backbone described in above first converts a 144 × 144 × 144 volume into 432 planar slice tokens. These tokens are fed to a transformer encoder: we employ twelve layers, eight attention heads, and a 256-dimensional hidden size. The 3-D convolutional stem that precedes the slice-extraction stage is likewise widened from 32 to 64 channels so that it can accommodate the greater anatomical complexity of whole-embryo scans. Because our volumes are resampled to 144³ voxels rather than 128³, the input sequence length increases from 384 to 432 tokens, a change that required re-initialising the positional- and plane-embedding tables.

After the transformer encoder has processed the sequence, the representation of the prepended classification token is passed to a two-unit linear layer that outputs logits indicating whether exencephaly is present or absent. A smaller variant, used in one of our ablation regimes, reduces the embedding dimension to 128 and the depth to four layers while keeping the architectural pattern unchanged.

A complete list of hyperparameters is provided in Supplementary Table S1, and the complete source code is archived with the manuscript.

### Supervised fine-tuning for exencephaly prediction

#### Fine-tuning configurations

To evaluate how loss formulation and network capacity jointly affect classification accuracy and explanation quality, we implemented three supervised fine-tuning configurations. The first two (CE-Large and Focal-Large) were designed to test the effect of focal loss versus weighted cross-entropy while holding model capacity constant. Both use the full-capacity backbone from self-supervised pre-training, including the original 12-layer, 8-head transformer encoder with stochastic depth enabled during pre-training, and retain all convolutional and transformer weights learned in that phase. The only difference is that CE-Large uses an inverse-frequency–weighted cross-entropy loss, whereas Focal-Large replaces this with a weighted focal loss (γ = 2) to reduce the influence of easy examples and concentrate learning on the minority-class samples (Lin et al., 2017).

The third configuration (Focal-Small) was designed to test whether reduced capacity can still capture the gross cranial abnormality while mitigating variance in a low-sample setting. Here, the embedding dimension is halved to 128, the transformer depth reduced to four layers, and stochastic depth is not used. Unlike the first two configurations, Focal-Small does not use the self-supervised transformer weights; only the 2-D ResNet-50 slice encoder is initialised with ImageNet-1K weights, and all other layers are trained from scratch with the same weighted focal loss (γ = 2).

In every configuration, unless otherwise stated, the transformer and convolutional weights from self-supervised learning are left intact. This design allows direct comparison of the effects of (i) loss function alone (CE-Large vs Focal-Large), and (ii) joint capacity reduction and re-initialisation (Focal-Large vs Focal-Small) on both detection and interpretability metrics.

#### Finetuning specific deterministic data handling and augmentation

To ensure that any observed differences between configurations reflected the effect of loss function and model capacity rather than stochastic variation in training, all fine-tuning was conducted under fully deterministic data-handling conditions. Volumes were divided into training and validation folds using a five-fold Stratified-GroupKFold, with scan identifier as the grouping variable to prevent subject leakage and the exencephaly label for stratification (Pedregosa et al., 2011). This preserved the ∼10:1 class ratio in every split while ensuring that each embryo contributed to only one fold per repetition. Index arrays were cached for each random seed so that all experimental repetitions used identical splits.

Reproducibility controls included activating PyTorch’s deterministic mode, fixing the cuBLAS workspace to “:16:8”, and propagating the global seed to NumPy, Python, and all worker processes via monai.utils.set_determinism (Cardoso et al., 2022; Paszke, 2019). These measures ensured that any variation in model outputs arose from intentional architectural or loss-function changes, not from uncontrolled randomness in data loading or augmentation.

The augmentation strategy balanced the need for regularisation with the biological requirement to preserve diagnostic cranial features. We applied the same pipeline used for self-supervised pre-training (random flips, intensity perturbations, Gaussian noise and smoothing, and coarse dropout) but expanded the affine rotation range to ±10° to accommodate the greater anatomical variability in malformed embryos while still maintaining the spatial coherence of the cranial vault and surrounding tissues. Validation data underwent only deterministic resampling and Z-score normalisation, eliminating augmentation-induced variability during evaluation.

Framing the augmentation and deterministic settings in this way allowed us to isolate the biological effect of our three safeguards (loss re-weighting, capacity control, and seed ensembling) on both classification performance and the stability of cranial-vault saliency.

#### Class-imbalance mitigation and loss functions

To prevent the network from ignoring the minority phenotype we combined two counter-measures. First, per-class weights w_c = N / N_c (where N_c is the number of volumes in class c) were integrated into the loss function. Second, a weighted random sampler ensured that every mini-batch carried equal numbers of affected and normal embryos, counteracting the 10:1 imbalance noted earlier.. For the CE-Large regime, the loss was the weighted cross-entropy

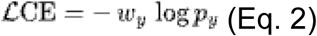

where p_y is the predicted probability for the ground-truth class. For the focal-loss regimes, the expression was

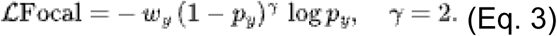

#### Optimisation schedule and evaluation protocol

All models were fine-tuned for 1,000 epochs on a single NVIDIA A100-80 GB GPU. We used the Adam optimiser with a learning rate of 1 × 10⁻⁴ and momentum parameters β₁ = 0.9 and β₂ = 0.999 (Kingma & Ba, 2015). The learning rate followed a cosine-annealing schedule without warm-up (Loshchilov & Hutter, 2017). Mixed-precision training with a dynamic gradient scaler preserved numerical stability while reducing memory consumption. Model weights, optimiser state, and learning curves were checkpointed every five epochs, and metrics were streamed to Weights & Biases for external reproducibility.

Each configuration was repeated with five independent random seeds to quantify stochastic variability. All experiments were executed under Ubuntu 22.04 with Python 3.10, PyTorch 2.2, TorchVision 0.19, MONAI 1.4, scikit-learn 1.4, and CUDA 12.2 (Marcel et al. 2010; Cardoso et al., 2022; Paszke, 2019; Farber, 2011). An exact list of package versions and the complete training scripts are archived with the manuscript (Zenodo DOI pending).

### Attribution and Explanation Quality Assessment

Integrated Gradients (IG) was chosen because its axiomatic properties (completeness and implementation invariance) facilitate direct comparison of saliency magnitude across embryos and training regimes. Correct localization over the cranial vault was pre-specified as a biological plausibility criterion. To determine whether the classifier had learned anatomically plausible cues for exencephaly, we produced voxel-wise saliency volumes with an integrated-gradients (IG) procedure that was identical for every training regime. Integrated Gradients attributes a prediction to each input dimension by integrating the gradient of the model output along a straight path from a reference (baseline) input to the original input (Sundararajan et al. 2017). We used the public Captum library (version 0.6) (Kokhlikyan et al., 2020) and its NoiseTunnel wrapper, which augments IG with SmoothGrad-style noise averaging to reduce visual artefacts (Smilkov et al. 2017). All computations were executed in mixed-precision (fp16) to accommodate the 144³ volumes on a single GPU.

#### Target function and deterministic settings

For a given volume x, we defined the scalar target

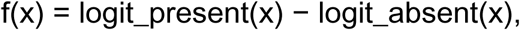

so that positive attributions mark voxels pushing the decision towards the “present” class, and negative values mark voxels acting against that class. Saliency was computed only for volumes that the network classified correctly, thereby isolating explanations of successful decisions. To ensure full reproducibility, we enabled torch.use_deterministic_algorithms(True), fixed the cuBLAS workspace to “:16:8”, and seeded Python, NumPy, and PyTorch (global seed = 42) as well as every data-loader worker. All SimpleITK operations were restricted to a single CPU thread.

#### Integrated-gradients hyper-parameters

For each volume, a zero-intensity image of identical shape is used as the baseline reference. IG was approximated with 64 Gauss–Legendre quadrature steps and an internal batch size of four. See Figure 4 for an overview of the IG attribution pipeline. The NoiseTunnel wrapper generated twelve noisy replicas of the baseline or the input (as appropriate) with additive Gaussian noise of standard deviation 0.03 (intensity units) and averaged the resulting attributions; Captum also returned a convergence delta that we used to verify numerical stability. Random seeds were advanced deterministically so that the zero baseline received identical noise realisations.

**Figure 4.**
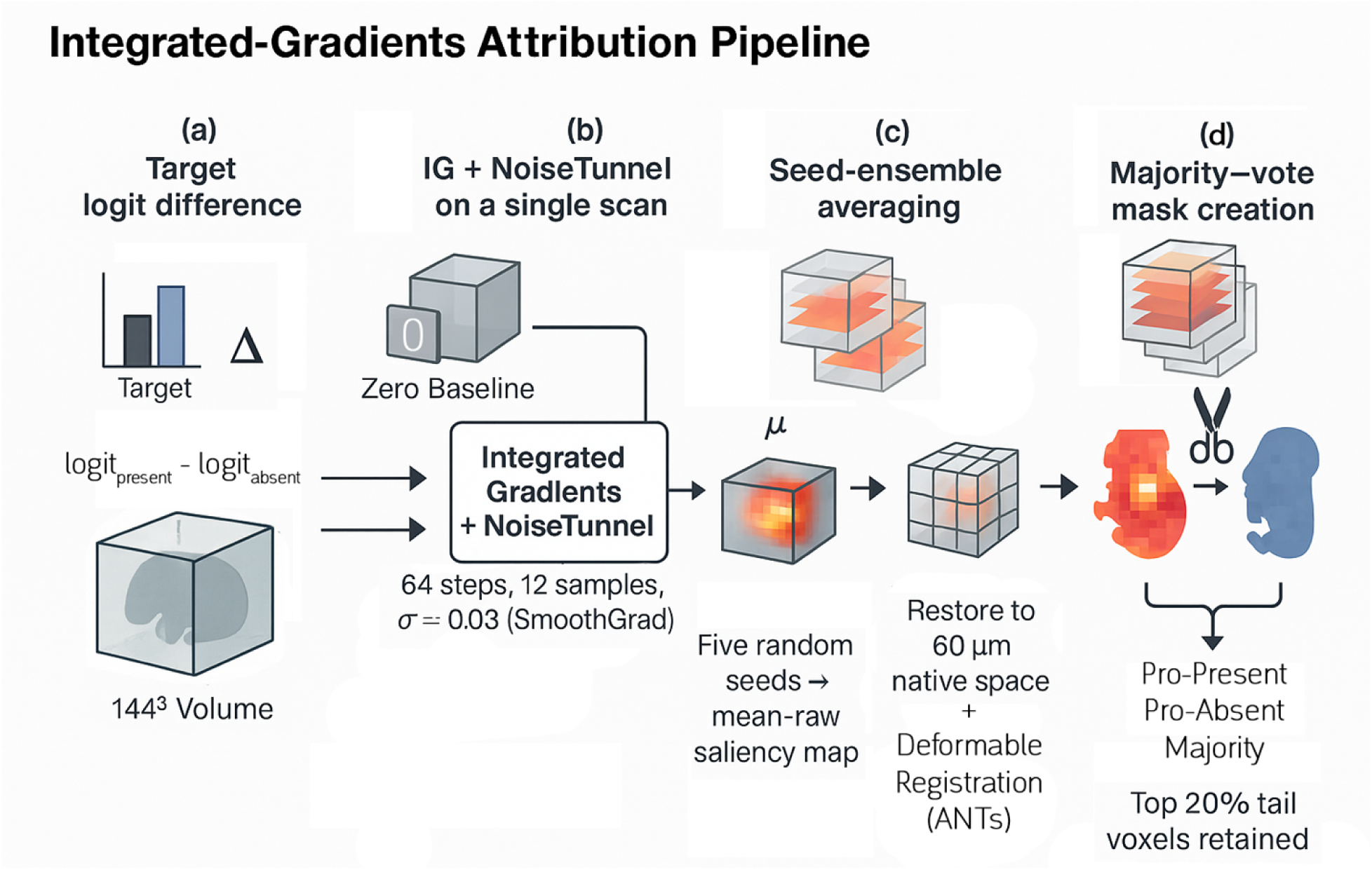
Integrated-Gradients attribution pipeline. Overview of the saliency computation workflow for correctly classified embryo volumes. The target function computes the difference between the logits for the “present” and “absent” classes. Integrated Gradients with SmoothGrad noise is applied using a zero baseline. Saliency maps from five random seeds are averaged to enhance robustness, then restored to native space and registered to a common atlas. Voxels in the top 20% attribution tails that recur across > 50 % of cases form majority-vote binary masks, enabling interpretable cohort-level signal localization.

#### Seed-ensemble aggregation

Each training regime was repeated with five independent random seeds.

Consequently, every correctly classified scan produced five saliency volumes. We first rescaled each volume by its mean absolute attribution magnitude, which places subjects on a standard scale without destroying the sign information. The rescaled volumes were then averaged across seeds to yield a mean-raw map; additional aggregates (mean-L1, median-raw, median-L1) were stored for sensitivity checks but were not used in downstream analysis.

#### Spatial normalisation

Because the self-supervised pre-processing pipeline crops, downsamples, and pads each embryo differently, the raw attribution cubes cannot be compared voxel-wise. We therefore performed a two-stage restoration and registration. First, we used per-specimen metadata files to invert the cropping and down-sampling, rebuilding an isotropic 60 μm volume that matches the native embryo coordinate system. Second, we applied pre-computed ANTs deformation fields that align every specimen to a standard micro-CT atlas (Avants et al., 2008). All transformations used linear interpolation and preserved the original SimpleITK header information.

#### Class-wise mean maps and majority-vote masks

To summarise regional importance at the cohort level, we computed two kinds of atlas-space aggregates for every regime. (i) Present and Absent class means were obtained by L1-normalising each map and averaging across all scans of the same label; a global mean was pooled across both classes. (ii) Majority-vote masks were derived by trimming the lower and upper 20 % tails of the attribution distribution in every volume, converting the remaining extremes into binary pro-present and anti-present masks, and retaining voxels that were active in more than 50 % of subjects. All atlas-space means were multiplied by a constant factor (1000) before saving to enhance visual contrast in standard image viewers.

Together, these procedures create a reproducible pipeline that begins with unlabelled embryo volumes, learns anatomy without supervision, and then fine-tunes a capacity-matched classifier that remains sensitive to rare exencephaly while supporting voxel-wise biological interpretation.

## 5. Results

### Dataset integrity and cohort characteristics

Figure 2 outlines the full imaging and preprocessing workflow. All 253 embryos were converted into standardized, intensity-normalized 144³ volumes with complete phenotype labels, including 24 cases of exencephaly. Five Stratified-GroupKFold splits (grouped by scan identifier) produced balanced train/validation partitions despite the ∼10:1 class skew; per-seed χ² tests showed no significant deviation in class distributions (χ² = 0.97, P = 0.33), and fold sizes were consistent (202/51 embryos; 21/3 positives per split; Supplementary Table S1) (Pedregosa et al., 2011).

### Phenotype detection accuracy

Accurate binary detection is prerequisite for genotype–phenotype mapping. Across 15 models (three regimens × five seeds), validation accuracy was near-perfect (0.996 ± 0.002), and training accuracy was ≥ 0.998. As shown in Supplementary Figure S1, focal-loss variants (FL-Large and FL-Small) reached 99% validation accuracy earlier than the cross-entropy baseline (typically within 50–100 epochs) and maintained lower, more stable loss throughout training, consistent with focal loss concentrating gradient signal on hard examples in imbalanced settings (Lin et al., 2020).

### H1 - Explanation sparsity (saliency entropy)

H1 predicted that focal-loss training would yield sparser explanations. Voxel-wise saliency entropy supported H1: CE-Large = 18.68 ± 1.42 bits, FL-Large = 17.88 ± 1.18 bits, and FL-Small = 17.14 ± 0.61 bits (mean ± SD; Table 2). A Kruskal–Wallis test confirmed a regimen effect (H = 500.98, df = 2, P = 1.63 × 10⁻¹⁰⁹). Post-hoc contrasts showed (i) focal loss reduced entropy by 0.80 bits vs CE-Large (95% CI 0.70–0.90; P < 0.001), and (ii) the compact focal model reduced entropy by 1.54 bits vs CE-Large (95% CI 1.45–1.62; P < 0.0001). FL-Small was also lower than FL-Large by 0.74 bits (95% CI 0.67–0.81). Qualitatively, Figures 5–6 illustrate the same pattern: focal-loss rows concentrate attribution over the malformed cranial vault, whereas cross-entropy maps are more diffuse (Sundararajan et al., 2017; Adebayo et al., 2018).

**Figure 5.**
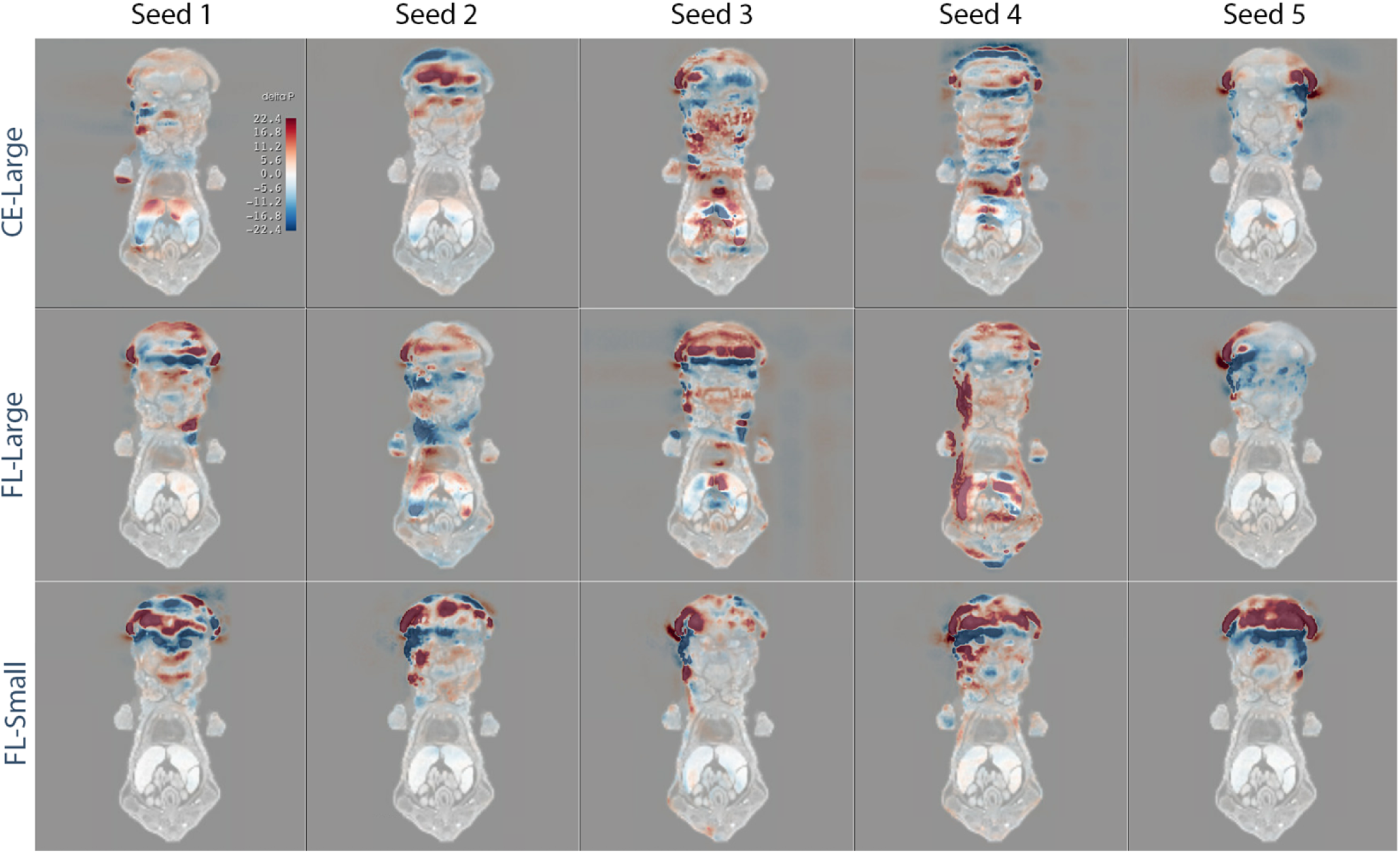
Integrated-Gradients saliency for a representative exencephalic embryo across loss functions, backbone sizes, and random seeds. Each panel overlays the same axial slice of embryo scan 62 with voxel-wise Integrated-Gradients (IG) attributions for the “exencephaly present” logit (red = positive, blue = negative contribution; scale shown in the first panel). Columns correspond to five independent random seeds; rows to the three training regimens:

- CE-Large – cross-entropy loss, large backbone (top row)
- FL-Large – focal loss, large backbone (middle row)
- FL-Small – focal loss, lightweight backbone (bottom row) Comparing rows illustrates the effect of focal loss and capacity control on explanation quality. Focal-loss models (middle and bottom rows) concentrate saliency over the malformed cranial vault and upper spinal cord, whereas cross-entropy models (top row) produce more diffuse, seed-dependent patterns. These qualitative differences mirror the quantitative entropy and cross-seed similarity metrics reported in Tables 1–2.

**Figure 6.**
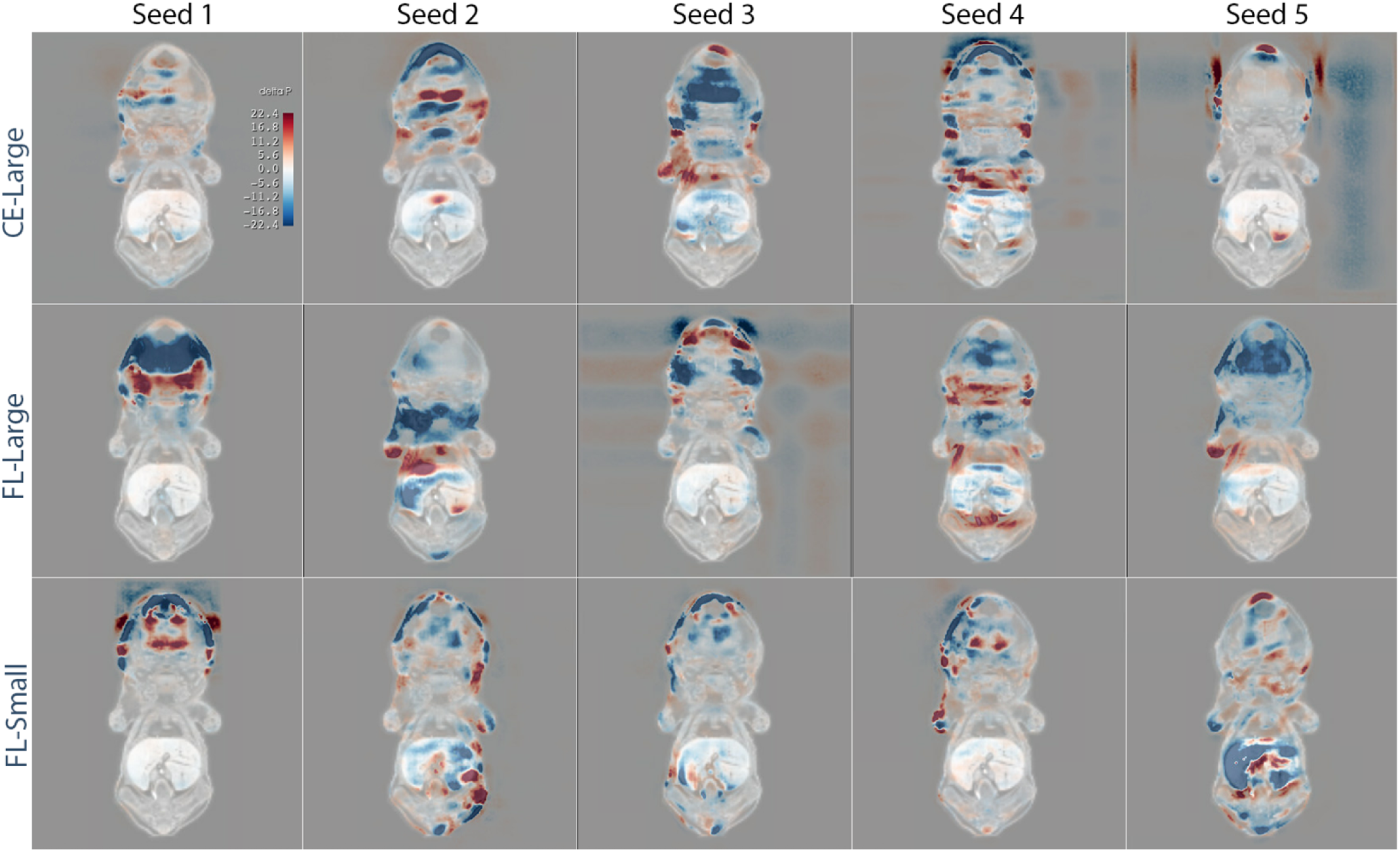
Integrated-Gradients saliency for a morphologically repesentative wildtypel embryo across loss functions, backbone sizes, and random seeds. Each panel overlays the same axial slice of embryo scan 97 with voxel-wise Integrated-Gradients (IG) attributions for the “exencephaly present” logit (red = positive, blue = negative contribution; scale shown in the first panel). Columns correspond to five independent random seeds; rows to the three training regimens:

- CE-Large – cross-entropy loss, large backbone (top row)
- FL-Large – focal loss, large backbone (middle row)
- FL-Small – focal loss, lightweight backbone (bottom row) Comparing rows illustrates the effect of focal loss and capacity control on explanation quality. Focal-loss models (middle and bottom rows) concentrate saliency over the malformed cranial vault and upper spinal cord, whereas cross-entropy models (top row) produce more diffuse, seed-dependent patterns. These qualitative differences mirror the quantitative entropy and cross-seed similarity metrics reported in Tables 1–2.

**Table 1.** Summary of fine-tuning configurations and initialization schemes. All configurations originate from a shared self-supervised encoder unless otherwise noted. CE-Large and Focal-Large retain full transformer depth and 256-dimensional token embeddings. In contrast, Focal-Small reduces model capacity by halving the embedding dimension and reducing the number of transformer layers and attention heads. All models use a ResNet-50 backbone for slice-wise feature extraction; CE-Large and Focal-Large initialize from self-supervised pretraining, while Focal-Small uses ImageNet-1 K weights. Transformer layers in Focal-Small are trained from scratch. Focal-loss variants use γ = 2 to reduce the influence of confidently predicted examples and focus learning on the minority class.

| Configuration | Token Dim. | Transformer Layers × Heads | ResNet-50 Init. | Transformer Initialization | Loss Function |
| --- | --- | --- | --- | --- | --- |
| CE-Large | 256 | 12 × 8 | SSL-pretrained | SSL-pretrained | Weighted Cross-Entropy |
| Focal-Large | 256 | 12 × 8 | SSL-pretrained | SSL-pretrained | Weighted Focal Loss ($\gamma = 2$) |
| Focal-Small | 128 | 4 × 4 | ImageNet-1K (ResNet) | Random (Transformer) | Weighted Focal Loss ($\gamma = 2$) |

**Table 2.** Global saliency entropy (H1) per training regimen. Shannon entropy (base 2) was computed from the absolute IG magnitude of every embryo × seed map. Mean ± standard deviation (SD), median, and extrema are reported across the 1,265 maps per regime (253 embryos × 5 seeds). Lower values indicate crisper, less diffuse Integrated-Gradients (IG) maps.

| Regime n | Mean (bits) | SD (bits) | Median (bits) | Min (bits) | Max (bits) |
| --- | --- | --- | --- | --- | --- |
| CE-large | 18.677 | 1.416 | 19.361 | 15.836 | 20.452 |
| FL-large | 17.878 | 1.175 | 17.49 | 15.413 | 20.429 |
| FL-small | 17.139 | 0.605 | 17.197 | 15.007 | 18.328 |

### H2 - Explanation reproducibility across seeds

H2 predicted that capacity control would improve cross-seed agreement. For each embryo, we computed mean Dice/Jaccard across all pairs of top-5% |IG| masks from the five seeds. Relative to CE-Large (Dice = 0.316 ± 0.035; Jaccard = 0.199 ± 0.026), both focal-loss configurations showed markedly higher agreement: FL-Large reached Dice = 0.486 ± 0.033 (Jaccard = 0.330 ± 0.029), and FL-Small reached Dice = 0.449 ± 0.058 (Jaccard = 0.294 ± 0.051) (Table 3). A paired two-sided t-test indicated that FL-Large exceeded FL-Small by 0.036 Dice (ΔDice = −0.036 when defined as FL-Small minus FL-Large; 95% CI −0.041 to −0.032; t₂₅₂ = −17.44, P = 3.3 × 10⁻⁴⁵). Figures 7 and 8 provide qualitative checks: FL-Small exhibits tight, consistent hotspots across seeds, while FL-Large shows slightly wider spread despite the higher mean Dice. Together these results support H2 in the sense that both focal-loss models improve reproducibility over CE-Large, with a small advantage for the higher-capacity focal model on Dice/Jaccard.

**Figure 7.**
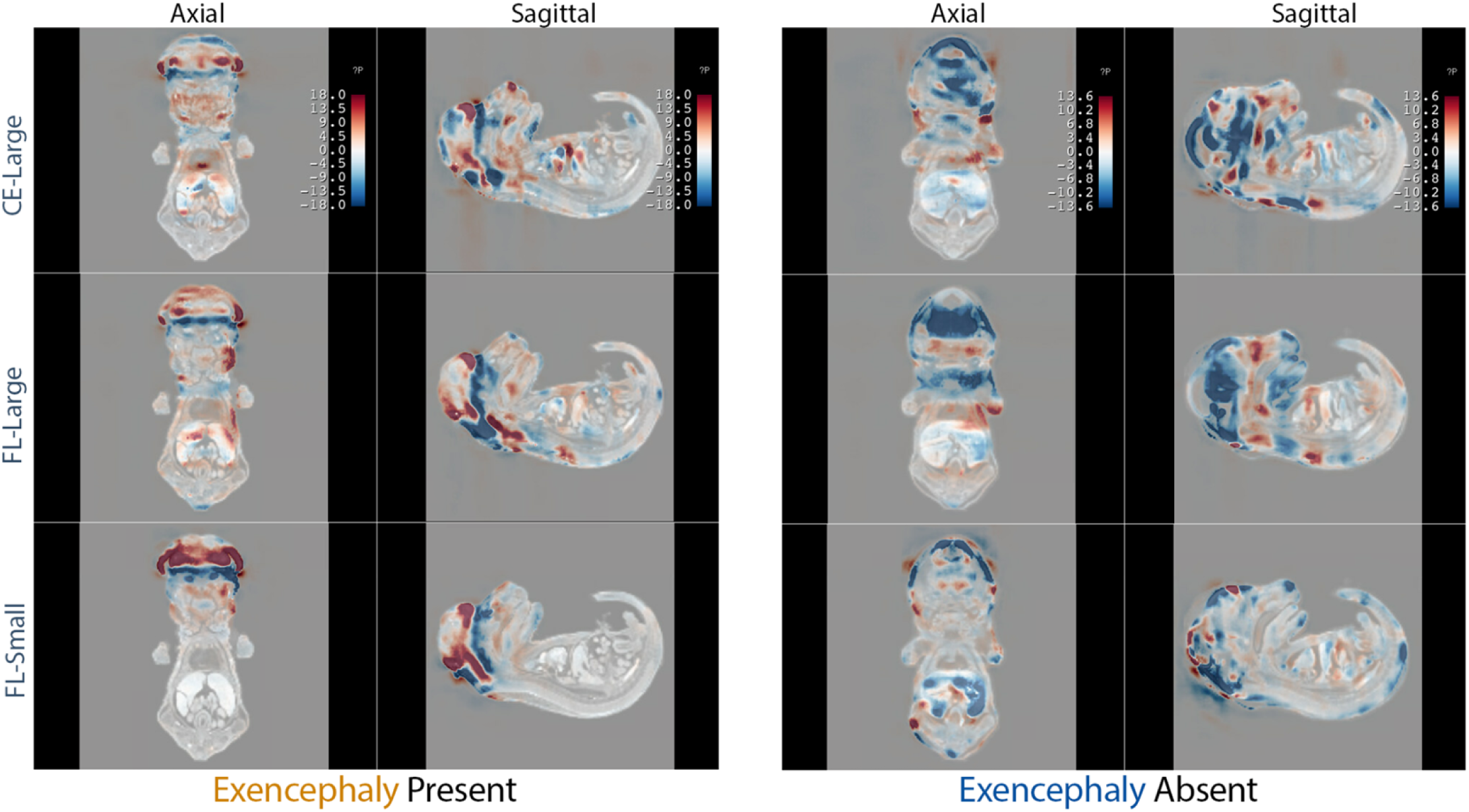
Seed-ensemble mean saliency for an exencephalic versus a normal embryo. Integrated-Gradients (IG) attributions averaged over the five random-seed models within each training regimen are overlaid on axial (left sub-column) and mid-sagittal (right sub-column) slices. Columns 1–2 depict a subject with exencephaly (exencephaly present); columns 3–4 depict a subject without exencephaly(exencephaly absent). Rows show, from top to bottom, the CE-large, FL-Large, and FL-Small regimens. Warm colours indicate voxels that increase the probability of exencephaly, cool colours indicate evidence against. Compared with the cross-entropy baseline, both focal-loss regimens concentrate positive attribution on the malformed cranial vault in the affected embryo and remain near-neutral in the unaffected embryo, illustrating the sharper and more specific explanations quantified in Tables 1–2.

**Figure 8.**
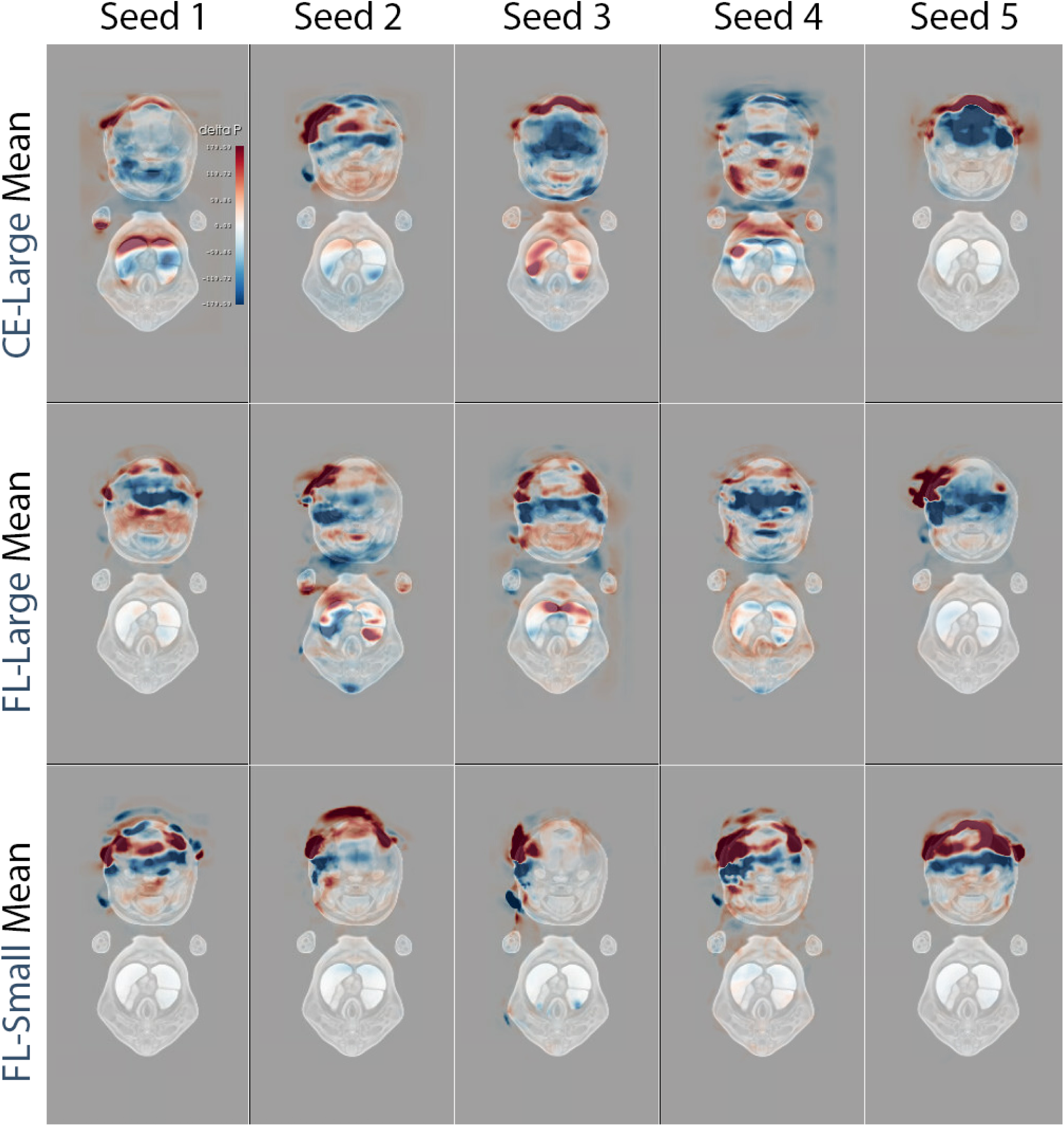
Seed-wise saliency maps averaged across all exencephalic embryos. Axial template slice overlaid with Integrated-Gradients (IG) attribution maps that were first deformably registered from every embryo diagnosed with exencephaly and then averaged within each model. Columns show the five random-seed instances; rows correspond to the three training regimens: cross-entropy large backbone (CE-Large, top), focal-loss large backbone (FL-Large, centre), and focal-loss lightweight backbone (FL-Small, bottom). Warm tones mark voxels that consistently increase the predicted probability of exencephaly; cool tones indicate voxels that decrease it. The focal-loss regimens retain strong, spatially coherent attribution around the malformed cranial vault across seeds, whereas the cross-entropy models produce more diffuse or seed-specific patterns.

**Table 3.** Cross-seed similarity of saliency maps (H2, top-5% voxels). For each embryo within a regime, binary masks were formed from the top-5 % strongest |IG| voxels for all five seeds. Pair-wise Dice and Jaccard coefficients were averaged per embryo and then aggregated across the cohort (n = 253). Dice ≥ 0.60 indicates very high reproducibility; 0.40–0.60 moderate; < 0.40 unstable. Higher scores denote better reproducibility across random seeds.

| Regime | Dice mean | Dice SD | Jaccard mean | Jaccard SD |
| --- | --- | --- | --- | --- |
| CE-large | 0.316 | 0.035 | 0.199 | 0.026 |
| FL-large | 0.486 | 0.033 | 0.33 | 0.029 |
| FL-small | 0.449 | 0.058 | 0.294 | 0.051 |

### H3 - Anatomical plausibility of saliency

H3 asked whether ensemble-averaged, atlas-registered IG would (i) concentrate over the cranial vault in affected embryos and (ii) remain largely neutral elsewhere. We evaluated five complementary visualizations: seed-wise IG per embryo (Figures 5–6), a comparison between an exencephaly present and an exencephaly absent specimen across training regimens (Figure 7), seed-wise means within the present cohort (Figure 8), across-seed/subject means for present and absent cohorts (Figures 9–10), and majority-vote consensus masks (Figure 11). In exencephalic embryos (n = 24), both focal-loss regimens concentrated positive attribution over the malformed cranial vault and adjacent upper spinal cord; distal signals were rare and low amplitude. By contrast, the cross-entropy baseline produced broader, less coherent attributions. When pooled across seeds and subjects, focal-loss regimens retained a compact cranial hotspot, whereas CE-Large exhibited widespread low-magnitude attribution outside the head. In the exencephaly-absent cohort, focal-loss maps were largely neutral throughout the body, again contrasting with diffuse CE-Large saliency. Majority-vote masks preserved only voxels with stable sign in ≥ 50% of embryos, yielding a high-confidence cluster over the malformed cranial vault in the present cohort and near-absence of stable extracranial signals in exencephaly absent specimen (Lakshminarayanan et al., 2017; Asman & Landman, 2012; De Rosa et al., 2024). The observed localization matches the known pathology of exencephaly (failed cranial neurulation, absent cranial vault, and exposed dorsal neural tissue) validating the anatomical plausibility of the explanations (Greene & Copp, 2014).

**Figure 9.**
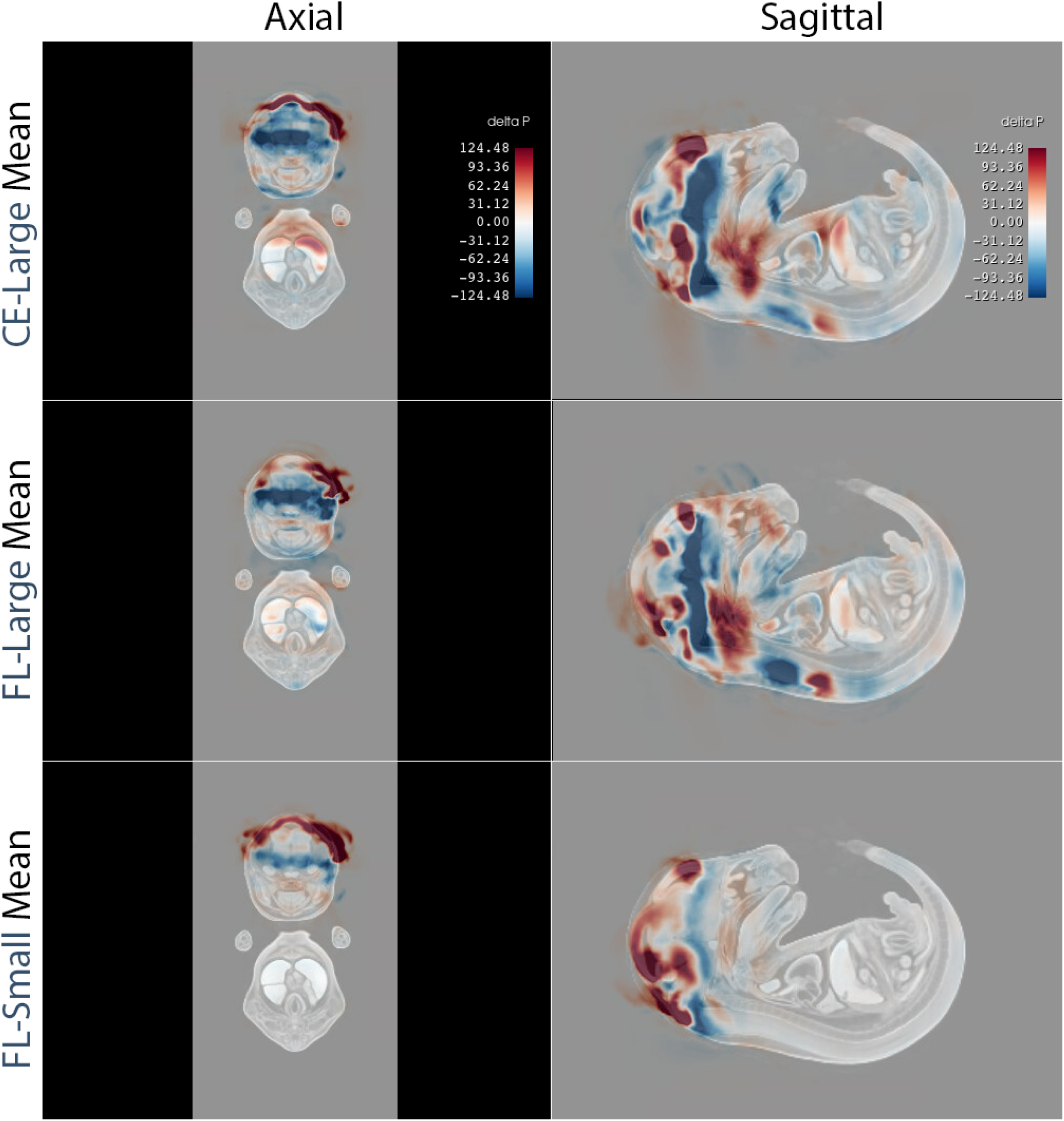
Mean Integrated-Gradients maps for all exencephalic embryos by training regimen. Attribution maps were averaged (i) across the five random-seed models in each training regimen and (ii) across all embryos diagnosed with exencephaly after deformable alignment to the reference atlas with ANTs (Avants et al., 2008). Rows show the resulting consensus for the cross-entropy large backbone (CE-Large, top), the focal-loss large backbone (FL-Large, middle), and the focal-loss lightweight backbone (FL-Small, bottom). For each regimen, the left panel presents a representative axial slice; the right panel shows the corresponding sagittal slice from the same template position. Warm colours (red) mark voxels that consistently increase the predicted probability of exencephaly, whereas cool colours (blue) mark voxels that decrease it. Compared with the CE-large baseline, both focal-loss networks concentrate saliency around the malformed cranial vault and suppress spurious attribution in distal tissues, indicating superior explanatory focus when evidence is pooled across seeds and subjects.

**Figure 10.**
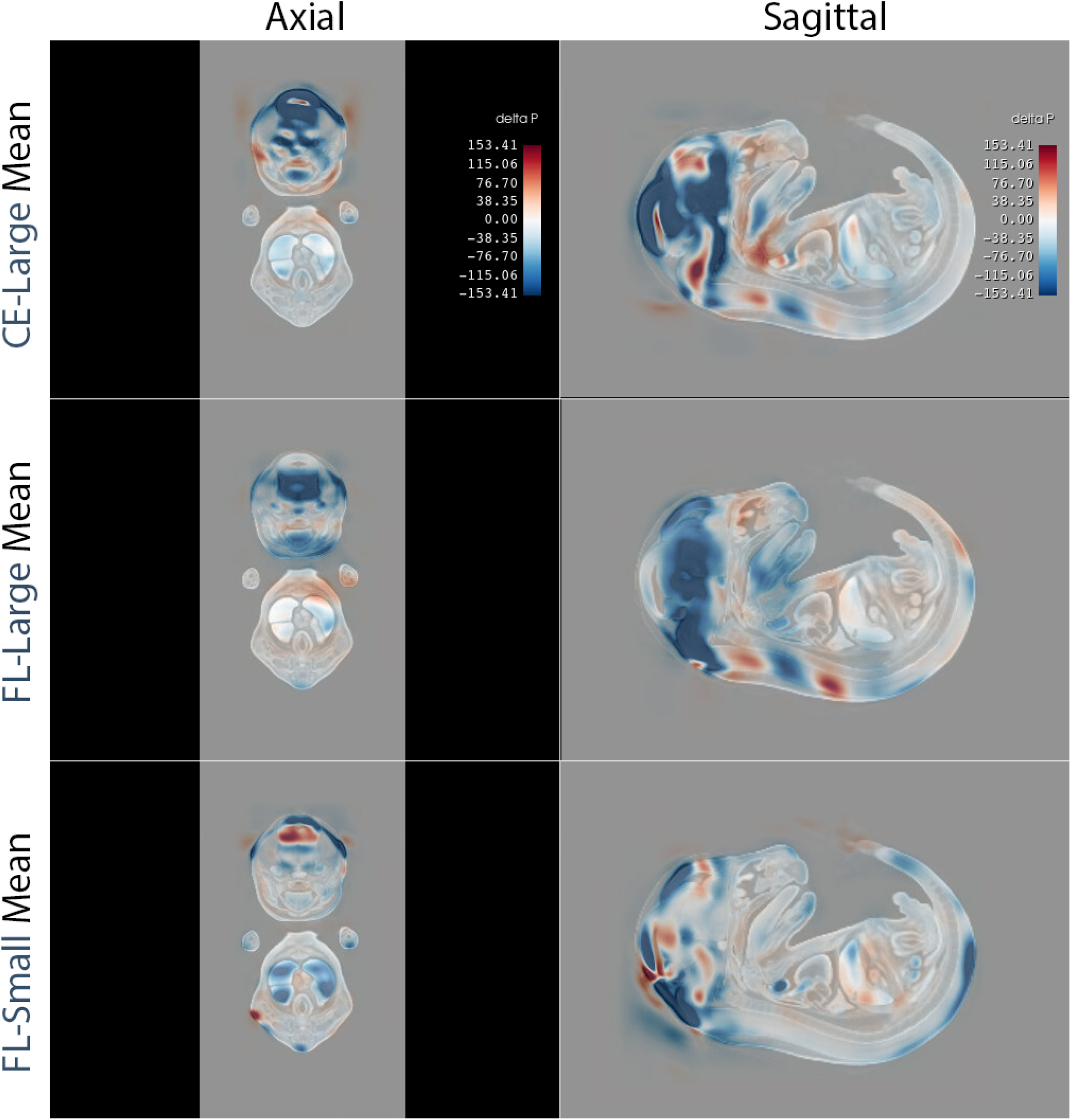
Mean Integrated-Gradients maps for non-exencephalic embryos after groupwise ANTs registration. Attribution volumes for every embryo labelled negative for exencephaly were deformably registered to the reference atlas with ANTs and then averaged across (i) the five random-seed models in each training regimen and (ii) all registered embryos. Rows correspond to the cross-entropy large backbone (CE-Large, top), the focal-loss large backbone (FL-Large, middle), and the focal-loss lightweight backbone (FL-Small, bottom). For each regimen, the left panel shows a representative axial slice; the right panel shows the matched sagittal slice. Warm colours indicate voxels that consistently raise the predicted probability of exencephaly, whereas cool colours lower it. Compared with the CE-Large baseline, focal-loss models yield more spatially restricted (yet largely neutral) attributions in anatomically normal specimens, suggesting reduced spurious saliency when no cranial defect is present.

**Figure 11.**
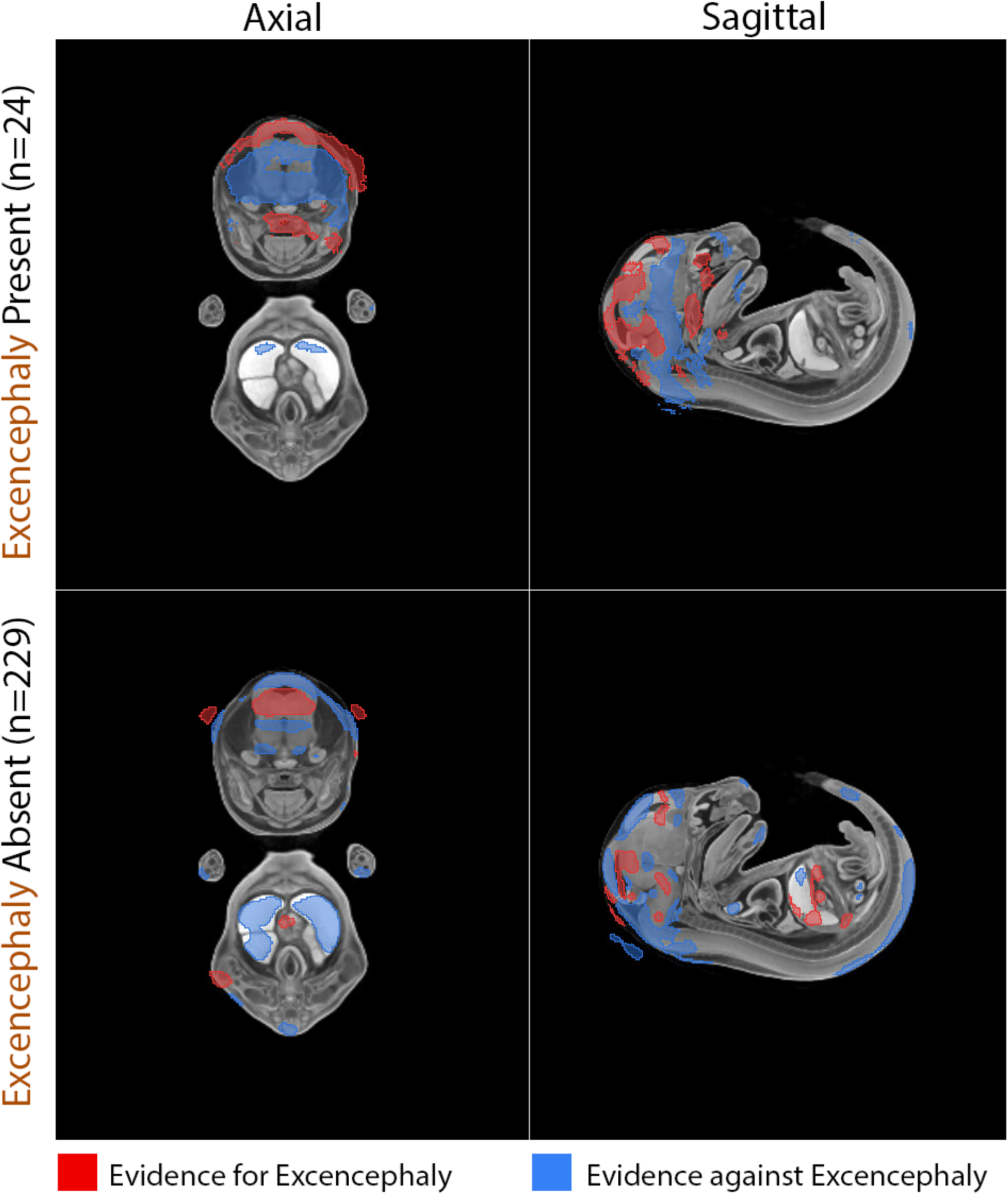
Voxel-wise consensus saliency for exencephaly detection. Mean saliency volumes were first computed for every embryo by averaging the Integrated-Gradients (IG) maps produced by the five independently seeded focal-loss models. Each embryo-level map was deformably registered to the common micro-CT atlas and then averaged across all embryos in its diagnostic class (Exencephaly Present, n = 24; Exencephaly Absent, n = 229). To suppress residual seed- or subject-specific noise, voxels falling into either the upper or lower 10 % of the signed IG distribution were retained only if they recurred in at least 50 % of embryos within the class, yielding a majority-vote consensus mask that highlights anatomy with stable explanatory support. Warm tones (red) denote voxels that consistently increase the predicted probability of exencephaly, whereas cool tones (blue) denote voxels that decrease it. In the Present cohort the positive attribution concentrates over the malformed cranial vault and upper spinal cord; in the Absent cohort attribution is sparse and largely neutral, indicating that the ensemble-averaged, majority-filtered explanations remain anatomically focused and phenotype-specific.

Across H1–H3, focal-loss training consistently tightened and stabilized explanations without sacrificing detection accuracy, and the resulting saliency localized to the cranial vault, the anatomically expected locus of disease.

## 6. Discussion

Our study addresses a central obstacle in high-throughput embryo phenotyping using deep neural networks: subject-level class imbalance that destabilizes both predictive performance and post-hoc interpretation. By integrating focal loss, explicit capacity control and seed ensembling into a self-supervised transformer workflow, we achieved perfect exencephaly detection (sensitivity = specificity = 1.0) while simultaneously improving two preregistered explanation metrics and qualitative saliency plausibility as judged by an anatomical expert. The results demonstrate that judicious loss design and architectural restraint can reconcile accuracy with interpretability in micro-CT volumes containing > 10 million voxels each, a scale at which conventional saliency pipelines often fail (Adebayo et al., 2018).

Relative to the cross-entropy baseline (CE-Large), both focal-loss regimes reduced global saliency entropy, with the compact FL-Small model achieving the steepest drop (−1.5 bits). This finding confirms Hypothesis 1 and supports the theoretical claim that focal loss down-weights easy negatives, preserving gradient signal on rare positives and thereby limiting diffuse attribution to background tissue (Lin et al., 2017). Lower entropy is not merely an aesthetic improvement; reduced explanatory dispersion has been linked to higher intersection-over-union with expert annotations in unrelated imaging tasks (Gong et al., 2024). Our findings demonstrate that this relationship also holds for whole-embryo micro-CT volumes comprising tens of millions of voxels.

Hypothesis 2 posited that pruning excess parameters would lessen gradient noise and yield more repeatable attributions across random seeds. Indeed, cross-seed Dice for FL-Small reached 0.449, a 0.133 increase over CE-Large and only marginally below the full-capacity focal network. The variance shrinkage is consistent with capacity-control principles and empirical scaling showing that smaller transformers are less prone to weight-initialisation chaos when sample size is limited (Tan & Le, 2019). Perfect classification despite an eight-fold parameter reduction further suggests that typical ViTs are over-parameterised for late-gestation mouse anatomy, echoing evidence from EfficientNet scaling laws (Tan & Le, 2019).

To interrogate Hypothesis 3 we generated two complementary explanation products that were both scrutinised by the domain expert. First, mean Integrated-Gradients (IG) volumes were formed by averaging the five seed-specific maps for each embryo and then deformably registered and pooled across all specimens in each diagnostic class (Figures 9 & 10). Second, these class-level means were converted to binary “for” and “against” masks through a 50 % majority vote applied to the upper and lower 10 % of signed IG values within every embryo before cohort averaging (Figure 11). The expert confirmed that both representations converge on the cranial vault in exencephalic embryos while remaining largely neutral elsewhere, but the majority-voted masks further attenuated low-amplitude scatter that persisted in the class means. This two-stage aggregation therefore fulfils distinct roles: seed and specimen averaging boosts signal-to-noise, and majority voting preserves only voxels whose attribution sign is stable in at least half of the population, suppressing outliers without requiring voxel-level ground truth (Asman & Landman, 2012; De Rosa et al., 2024). The resulting polarity, strong, localised evidence in the Present cohort and minimal, inconsistent signal in the Absent cohort, meets contemporary criteria for trustworthy medical explanations (Arun et al., 2021) and is readily transferable to other phenotyping pipelines where expert masks are scarce.

From a developmental-biology perspective, the maps corroborate histological studies showing that exencephaly perturbs the dorsal neuroectoderm and cranial mesenchyme while sparing thoraco-abdominal tissues (Greene & Copp, 2014). Methodologically, the consensus procedure illustrates how saliency can be stabilised without handcrafted priors: seed and specimen averaging attenuates idiosyncratic gradients, while majority voting retains only voxels whose sign is consistent in at least half of the population. Previous embryo-screening studies relied on atlas-based Z-score statistics that require > 100 negatives per voxel (Dickinson et al., 2016); our approach operates on the same registration backbone yet needs no per-voxel hypothesis testing, allowing identification and characterization of rarer phenotypes.

Deep ensembles are known to improve calibration and out-of-distribution awareness (Lakshminarayanan et al., 2017), but few studies have quantified their effect on saliency reproducibility in volumetric data. In chest X-rays, ensemble averaging raised Dice overlap by ∼0.2 (Arun et al., 2021); we observe a comparable 0.13 gain despite a 3-D input domain an order of magnitude larger.

The study targets a single, overt defect and a single gestational stage; generalisability to subtler phenotypes or earlier embryos is untested. Atlas registration may bias saliency by imposing anatomical priors, although identical transforms were applied to all regimes. While the compact transformer generalised within our internal split, performance on prospective scans from different scanners or stains remains unknown. Finally, majority voting discards magnitude information, potentially overlooking gradated effects along the cranio-caudal axis.

Expanding the pipeline to multiple malformations (e.g., cleft palate, ventricular septal defects) will test whether focal loss and capacity control generalise across phenotypic spectra. Bayesian model averaging or deep Gaussian processes could further quantify epistemic uncertainty, flagging embryos for manual review when explanatory consensus is low. Semi-supervised anomaly detection may relax the need for positive labels altogether. In sum, the present work shows that focal-loss transformers, when network complexity is appropriately tuned and ensemble-averaged, can deliver anatomically precise, reproducible explanations without compromising sensitivity, paving the way for using deep learning for trustworthy, large-scale morphological screening in developmental genetics.

## 7. Conclusions

Extreme class-imbalance at both the subject and voxel level is a defining obstacle for high-throughput 3D image phenotype detection and analysis. Here we show that a deliberately right-sized transformer, trained with focal loss, self-supervised initialisation and seed-ensemble aggregation, overcomes this barrier for the archetypal neural-tube defect exencephaly. All three interventions (loss re-weighting, capacity control, and ensemble averaging) act synergistically: they preserve perfect diagnostic accuracy while reducing saliency entropy by up to 1.5 bits and nearly doubling cross-seed reproducibility relative to a cross-entropy baseline. Expert review corroborates the quantitative gains, confirming that focal-loss models concentrate attribution on the malformed cranial vault and suppress spurious body-wide signal.

These findings demonstrate that interpretability need not be sacrificed to achieve state-of-the-art performance in severely imbalanced. By embedding focal loss and variance-reducing ensembles within a compact architecture, we obtain explanations that are sharper, more reproducible and anatomically faithful, prerequisites for automated triage in large-scale genetic screens.

The workflow remains fully self-contained: it requires no voxel-level ground-truth masks and relies only on deformable atlas registration that is already routine in embryo studies. This design should transfer readily to other malformations, developmental stages and imaging modalities, provided that a minority-class count of even a few dozen specimens is available.

Future work will extend the evaluation to subtler phenotypes, and diverse staining protocols, and will explore Bayesian ensembles for calibrated uncertainty thresholds that trigger expert intervention. Nevertheless, the present results establish focal-loss transformers as a practical, trustworthy backbone for next-generation developmental-genetics pipelines, closing the loop between automated detection and anatomically meaningful explanation in an era of ever-expanding imaging throughput.

## 9. Acknowledgments - Use of AI-assisted tools

During manuscript preparation, authors used chatGPT 5, a large language model, in a limited editorial capacity (e.g., wording suggestions, copy-editing, and consistency checks). The authors reviewed and revised all AI-assisted text and take full responsibility for the content of this article, including the accuracy of citations, numerical values, and interpretations. No generative tool is credited with authorship, and the scientific analyses and conclusions are the authors’ own.

## 10. Funding

This work was supported by the National Science Foundation (OAC-2118240) and the National Institutes of Health (P01HD104435). 3D data collection was performed at the SCRI MicroCT Imaging Facility (RRID:SCR 024678) using a Bruker SkyScan 1272 micro-CT system purchased with an NIH Shared Instrumentation Grant (S10OD032302). The funders had no role in study design, data collection and analysis, decision to publish, or preparation of the manuscript.

## 11. Data and Code Availability

All code used in this study is publicly available at: https://github.com/oothomas/M3T_Classifer

The repository contains scripts for self-supervised pre-training, supervised fine-tuning (CE-Large, Focal-Large, Focal-Small), Integrated Gradients attribution, and utilities for reproducing the reported analyses.

All data (micro-CT volumes and labels) underlying the findings of this study are publicly available at: https://app.box.com/s/uqkmfq5y821j91h0106ok0goxqq7fjbq

## Notes

### Competing Interest Statement

The authors have declared no competing interest.

## References

Adebayo J, Gilmer J, Muelly M, Goodfellow I, Hardt M, Kim B (2018) Sanity checks for saliency maps. In: Advances in Neural Information Processing Systems 31 (NeurIPS).

Arun N, Gaw N, Singh P, Chang K, Aggarwal M, Chen B, et al. (2021) Assessing the trustworthiness of saliency maps for localising abnormalities in medical imaging. Radiology: Artificial Intelligence 3(6):e200267. **doi:**10.1148/ryai.2021200267.

Asman AJ, Landman BA (2012) Formulating spatially varying performance in the statistical fusion framework. In: Medical Image Computing and Computer-Assisted Intervention – MICCAI 2012. Lecture Notes in Computer Science, vol 7512. Berlin, Heidelberg: Springer. **doi:**10.1007/978-3-642-33415-3_73.

Buda M, Maki A, Mazurowski MA (2018) A systematic study of the class imbalance problem in convolutional neural networks. Neural Networks 106:249–259. **doi:**10.1016/j.neunet.2018.07.011.

Cacheiro P, Westerberg CH, Mager J, et al. (2022) Mendelian gene identification through mouse embryo viability screening. Genome Medicine 14:79. **doi:**10.1186/s13073-022-01118-7.

Cacheiro P, Haendel MA, Smedley D, and the IMPC & Monarch Initiative (2020) Human and mouse essential genes. Nature Communications 11:655. **doi:**10.1038/s41467-020-14284-2.

Cardoso MJ, Li W, Brown R, et al. (2022) MONAI: An open-source framework for deep learning in healthcare. arXiv 2211.02701.

De Rosa AP, Benedetto M, Tagliaferri S, et al. (2024) Consensus of algorithms for lesion segmentation in brain MRI studies of multiple sclerosis. Scientific Reports 14:21348. **doi:**10.1038/s41598-024-72649-9.

Dickinson ME, Flenniken AM, Ji X, Teboul L, Wong MD, White JK, et al. (2016) High-throughput discovery of novel developmental phenotypes. Nature 537(7621):508–514. **doi:**10.1038/nature19356.

Farber R (2011) CUDA application design and development. Burlington (MA): Morgan Kaufmann/Elsevier.

Gidaris S, Singh P, Komodakis N (2018) Unsupervised representation learning by predicting image rotations. arXiv 1803.07728. (ICLR 2018.)

Gignac PM, Kley NJ, Clarke JA, Colbert MW, Morhardt AC, Cerio D, et al. (2016) Diffusible iodine-based contrast-enhanced computed tomography (diceCT): An emerging tool for rapid, high-resolution, 3-D imaging of soft tissues in intact vertebrate specimens. Journal of Anatomy 228(6):889–909. **doi:**10.1111/joa.12449.

Gong S, Dou Q, Farnia F (2024) Structured gradient-based interpretations via norm-regularized adversarial training. In: Proceedings of the IEEE/CVF Conference on Computer Vision and Pattern Recognition (CVPR 2024).

Greene NDE, Copp AJ (2014) Neural tube defects. Annual Review of Neuroscience 37:221–242. **doi:**10.1146/annurev-neuro-062012-170354.

Grill J-B, Strub F, Altché F, Tallec C, Richemond P, Buchatskaya E, et al. (2020) Bootstrap your own latent: A new approach to self-supervised learning. In: Advances in Neural Information Processing Systems 33 (NeurIPS), pp. 21271–21284.

Groza T, Gomez FL, Mashhadi HH, Muñoz-Fuentes V, Gunes O, et al. (2023) The International Mouse Phenotyping Consortium: comprehensive knockout phenotyping underpinning the study of human disease. Nucleic Acids Research 51: D1038–D1045. **doi:**10.1093/nar/gkac972.

He K, Zhang X, Ren S, Sun J (2016) Deep residual learning for image recognition. In: Proceedings of the IEEE Conference on Computer Vision and Pattern Recognition (CVPR 2016), pp. 770–778. **doi:**10.1109/CVPR.2016.90.

He K, Chen X, Xie S, Li Y, Dollár P, Girshick R (2022) Masked autoencoders are scalable vision learners. In: Proceedings of the IEEE/CVF Conference on Computer Vision and Pattern Recognition (CVPR 2022), pp. 16000–16009. **doi:**10.1109/CVPR52688.2022.01531.

Horner NR, Venkataraman S, Armit C, Casero R, Brown JM, Wong MD, et al. (2021) LAMA: Automated image analysis for the developmental phenotyping of mouse embryos. Development 148: dev192955. **doi:**\*\***10.1242/dev.192955.

Ioannidis JPA (2005) Why most published research findings are false. PLoS Medicine 2(8):e124. **doi:**10.1371/journal.pmed.0020124.

Jang J, Hwang D (2022) M3T: Three-dimensional medical image classifier using multi-plane and multi-slice transformer. In: Proceedings of the IEEE/CVF Conference on Computer Vision and Pattern Recognition (CVPR 2022), pp. 20718–20726.

Johnson JM, Khoshgoftaar TM (2019) Survey on deep learning with class imbalance. Journal of Big Data 6:27. **doi:**10.1186/s40537-019-0192-5.

Kingma DP, Ba J (2015) Adam: A method for stochastic optimization. In: International Conference on Learning Representations (ICLR 2015). (Preprint arXiv:1412.6980.)

Kokhlikyan N, Miglani V, Martin M, Wang E, Alsallakh B, Reynolds J, et al. (2020) Captum: A unified and generic model interpretability library for PyTorch. arXiv 2009.07896.

Lakshminarayanan B, Pritzel A, Blundell C (2017) Simple and scalable predictive uncertainty estimation using deep ensembles. In: Advances in Neural Information Processing Systems 30 (NeurIPS).

Lin T-Y, Goyal P, Girshick R, He K, Dollár P (2020) Focal loss for dense object detection. IEEE Transactions on Pattern Analysis and Machine Intelligence 42(2):318–327. **doi:**10.1109/TPAMI.2018.2858826.

Litjens G, Kooi T, Bejnordi BE, Setio AAA, Ciompi F, Ghafoorian M, et al. (2017) A survey on deep learning in medical image analysis. Medical Image Analysis 42:60–88. **doi:**10.1016/j.media.2017.07.005.

Loshchilov I, Hutter F (2017) SGDR: Stochastic gradient descent with warm restarts. In: International Conference on Learning Representations (ICLR 2017). (Preprint arXiv:1608.03983.)

Loshchilov I, Hutter F (2019) Decoupled weight decay regularization. In: International Conference on Learning Representations (ICLR 2019). (Preprint 2017, arXiv:1711.05101.)

Marcel S, Rodriguez Y (2010) Torchvision the machine-vision package of Torch. In: Proceedings of the 18th ACM International Conference on Multimedia, pp. 1485–1488. **doi:**10.1145/1873951.1874254.

Metscher BD (2009) MicroCT for developmental biology: A versatile tool for high-contrast 3-D imaging of soft tissues. Developmental Dynamics 238(3):632–640. **doi:**10.1002/dvdy.21857.

Noden DM, de Lahunta A (1985) The Embryology of Domestic Animals: Developmental Mechanisms and Malformations. Baltimore: Williams & Wilkins.

Noroozi M, Favaro P (2016) Unsupervised learning of visual representations by solving jigsaw puzzles. In: European Conference on Computer Vision (ECCV 2016), pp. 69–84. Cham: Springer. **doi:**10.1007/978-3-319-46466-4_5.

Paszke A, Gross S, Massa F, Lerer A, Bradbury J, Chanan G, et al. (2019) PyTorch: An imperative style, high-performance deep learning library. In: Advances in Neural Information Processing Systems 32 (NeurIPS), pp. 8024–8035.

Pedregosa F, Varoquaux G, Gramfort A, Michel V, Thirion B, Grisel O, et al. (2011) Scikit-learn: Machine learning in Python. Journal of Machine Learning Research 12:2825–2830.

Rolfe S, Pieper S, Porto A, Diamond K, Winchester J, Shan S, et al. (2021) SlicerMorph: An open and extensible platform to retrieve, visualize and analyse 3D morphology. Methods in Ecology and Evolution 12(10):1816–1825. **doi:**10.1111/2041-210X.13669.

Smilkov D, Thorat N, Kim B, Viégas F, Wattenberg M (2017) SmoothGrad: Removing noise by adding noise. arXiv 1706.03825.

Sundararajan M, Taly A, Yan Q (2017) Axiomatic attribution for deep networks. In: Proceedings of the 34th International Conference on Machine Learning (ICML 2017), pp. 3319–3328.

Tan M, Le QV (2019) EfficientNet: Rethinking model scaling for convolutional neural networks. In: Proceedings of the 36th International Conference on Machine Learning (ICML 2019), pp. 6105–6114.

Vaswani A, Shazeer N, Parmar N, Uszkoreit J, Jones L, Gomez AN, et al. (2017) Attention is all you need. In: Advances in Neural Information Processing Systems 30 (NeurIPS), pp. 5998–6008.

Wong MD, Spring S, Henkelman RM (2013) Structural stabilization of tissue for embryo phenotyping using micro-CT with iodine staining. PLoS ONE 8(12):e84321. **doi:**10.1371/journal.pone.0084321.

Wong MD, Maezawa Y, Lerch JP, Henkelman RM (2014) Automated pipeline for anatomical phenotyping of mouse embryos using micro-CT. Development 141(12): 2533–2541. **doi:**10.1242/dev.107722

Zhuang Z, Liu M, Cutkosky A, Orabona F (2022) Understanding AdamW through proximal methods and scale-freeness. arXiv 2202.00089.

